# Root volatiles in plant-plant interactions II: Root terpenes from *Centaurea stoebe* modify *Taraxacum officinale* root chemistry and root herbivore growth

**DOI:** 10.1101/441790

**Authors:** Wei Huang, Valentin Gfeller, Matthias Erb

## Abstract

Volatile organic compounds (VOCs) emitted by plant roots can influence the germination and growth of neighboring plants. However, little is known about the effects of root VOCs on plant-herbivore interactions. The spotted knapeed (*Centaurea stoebe*) constitutively releases high amounts of sesquiterpenes into the rhizosphere. Here, we examine the impact of *C. stoebe* root VOCs on primary and secondary metabolites of sympatric *Taraxacum officinale* plants and the resulting plant-mediated effects on a generalist root herbivore, the white grub *Melolontha melolontha.* We show that exposure of *T. officinale* to *C. stoebe* root VOCs does not affect the accumulation of defensive secondary metabolites, but modulates carbohydrate and total protein levels in *T. officinale* roots. Furthermore, VOC exposure increases *M. melolontha* growth on *T. officinale* plants. Exposure of *T. officinale* to a major *C. stoebe* root VOC, the sesquiterpene (*E*)-β-caryophyllene, partially mimics the effect of the full root VOC blend on *M. melolontha* growth. Thus, releasing root VOCs can modify plant-herbivore interactions of neighboring plants. The release of VOCs to increase the susceptibility of other plants may be a form of plant offense.

## Introduction

Plants emit a variety of volatile organic compounds (VOCs) that can affect the behavior and performance of other organisms. VOCs induced by herbivory for instance can enhance defenses and resistance of neighboring plants (Arimura et al., 2000; Engelberth, Alborn, Schmelz, & Tumlinson, 2004; Frost, Mescher, Carlson & De Moraes, 2008, Erb et al., 2015; Karban, Yang, & Edwards, 2014; Pearse, Hughes, Shiojiri, Ishizaki, & Karban, 2013; Sugimoto et al., 2014). As the benefit for the emitter plant is unclear, this phenomenon is commonly regarded as a form of “eavesdropping” by the receiver rather than a form of communication (Heil & Karban, 2010). From the perspective of an emitter plant, it would seem advantageous to use VOCs to suppress rather than enhance defenses in neighbors (Heil & Karban, 2010). However, little is known about the capacity of VOCs to suppress defenses and enhance herbivore attack rates in neighboring plants. Broccoli plants were found to receive more oviposition by diamondback moths after exposure to VOCs from damaged conspecifics (Li & Blande, 2015). Furthermore, exposure to VOCs from damaged neighbors increases herbivore damage on blow-wives (*Achyrachaena mollis*) and charlock (*Sinapis arvensis*) (Pearse et al., 2012). Finally, GLV exposure suppresses several defense-related genes in coyote tobacco (*Nicotiana attenuata*) (Paschold, Halitschke, & Baldwin, 2006). Clearly, the capacity of VOCs to suppress rather than induce defenses requires more attention in order to understand how VOCs influence plant-herbivore interactions of neighboring plants (Erb, 2018b).

The majority of studies on the effects of VOCs on plant neighbors have focused on the phyllosphere. However, plants also release significant amounts of VOCs into the rhizosphere, which may affect plant defense and plant-herbivore interactions (Delory, Delaplace, Fauconnier, & du Jardin, 2016). Root VOCs can affect the germination and growth of neighboring plants (Ens, Bremner, French, & Korth, 2009; Jassbi, Zamanizadehnajari, & Baldwin, 2010) and the performance of herbivores (Hu et al., 2018a; Robert et al., 2012). Therefore, it is reasonable to assume that they may also affect plant-herbivore interactions of neighboring plants. Root exudates and mycelial networks have been shown to alter plant defenses and plant herbivore interactions in neighboring plants (Babikova et al., 2013; Dicke & Dijkman, 2001), but the specific role of root VOCs in plant-plant interaction has, to the best of our knowledge, not been addressed (Delory et al., 2016).

In this study, we explored the influence of root VOCs on the common dandelion (*Taraxacum officinale* agg.) and its interaction with the common cockchafer *Melolontha melolontha*. In grasslands across Europe, *T. officinale* is often attacked by larvae of *M. melolontha* (Coleoptera, Scarabaeidae) (Huber et al., 2016a), a highly polyphagous root feeder (Hauss & Schütte, 1976; Sukovata, Jaworski, Karolewski, & Kolk, 2015). Previous work found that the interaction between *T. officinale* and *M. melolontha* is modulated by the presence of sympatric plant species (Huang, Zwimpfer, Hervé, Bont, & Erb, 2018). Strong effects were for instance observed for *Centaurea stoebe,* a native herb that is invasive in the United States. *Melolontha melolontha* larvae grew significantly better on *T. officinale* plants in the presence of *C. stoebe,* an effect which was found to be mediated through changes in *T. officinale* susceptibility rather than direct effects of *C. stoebe* on the herbivore (Huang et al., 2018). In a companion paper, we describe that *C. stoebe* constitutively produces and releases significant amounts of sesquiterpenes into the rhizosphere (companion paper Gfeller et al., under review). Furthermore, we show that *C. stoebe* root VOCs have neutral to positive effects on the germination and growth of different neighboring species (companion paper Gfeller et al., under review). Based on these results, we hypothesized that *C. stoebe* root VOCs may play a role in increasing *T. officinale* susceptibility to *M. melolontha*. We tested this hypothesis by exposing *T. officinale* plants to root VOCs from *C. stoebe* and a major *C. stoebe* sesquiterpene and measuring changes in root primary and secondary metabolites and *M. melolontha* growth. This work provides evidence that root VOCs can influence plant-herbivore interactions on neighboring plants.

## Methods and Materials

### Study system

The study system consisted *T. officinale* (Genotype A34) as a receiver plant, *C. stoebe* as an emitter plant and *M. melolontha* as an herbivore of *T. officinale. Taraxacum officinale* seeds were obtained from greenhouse-grown A34 plants. *Centaurea stoebe* L. (diploid) seeds were obtained from a commercial vendor (UFA-SAMEN, Winterthur, Switzerland). *Melolontha melolontha* larvae were collected from an apple tree yard in Sion, Switzerland (46.21°N, 7.38°E). The larvae were reared on carrot slices under controlled condition (12°C, 60% humidity and constant darkness) for several weeks until the start of the experiments.

### Impact of C. stoebe root VOCs on the interaction between T. officinale and M. melolontha

To examine whether root VOCs emitted by *C. stoebe* affect the interaction between *T. officinale* and *M. melolontha*, *C. stoebe* and *T. officinale* plants were grown in pairs in an experimental setup that allowed only VOCs to diffuse from one plant to the other. Using this setup, we tested the effect of *C. stoebe* volatiles on *T. officinale* physiology (n=8) and on the growth of *M. melolontha* on *T. officinale* (n=16) as follows: Seeds of *T. officinale* and *C. stoebe* were germinated in the greenhouse at 50-70 % relative humidity, 16/8 h light/dark cycle, and 24 °C at day and 18°C at night. Ten days later, two seedlings of each species were transplanted into a mesh cage (12 × 9 × 10 cm, length × width × height) filled with a mixture of 1/3 landerde (Ricoter, Switzerland) and 2/3 seedling substrate (Klasmann-Deilmann, Switzerland). The mesh cage was made of geotex fleece (Windhager, Austria). Then, two mesh cages were put into a 2 L rectangular pot (18 × 12 × 10 cm, length × width × height). To reduce the interaction between focal and neighboring plants through root exudates, the mesh cages in each pot were separated by two plastic angles (0.8 cm width) and the pot was cut to produce a gap (12 × 0.5 cm, length × width) in the center of the bottom paralleling to the longest side of mesh cage. Finally, the gap in the top between two mesh cages was covered by a plastic sheet. A schematic drawing of the setup is shown in Fig. 1A. The setup is identical to the one used in the companion paper (companion paper Gfeller et al., under review). Seven weeks after transplantation, a pre-weighted *M. melolontha* larva was added into the mesh cage with focal plants. The larvae had been starved for three days prior to the experiment. After 18 days of infestation, the larvae were removed and re-weighted. Then, roots of focal plants were harvested, weighted and stored in −80 °C for further chemical analyses including soluble protein and sugars as well as the defensive metabolite sesquiterpene lactone taraxinic acid β-D glucopyranosyl ester (TA-G). Soluble protein was estimated using the Bradford method (Bradford, 1976). Soluble sugars including glucose, fructose and sucrose were measured as described by Velterop & Vos (2001) and Machado et al. (2013). TA-G was analyzed as described by Huber et al. (2015) and Bont et al. (2017). During the experiment, pots were watered daily. Care was taken not to overwater the plants to avoid leachate to cross the airgap between the inner mesh cages. The plant pairs were arranged randomly on a greenhouse table, with distances between pairs equal to distances within pairs. The positions of the pots on the table were re-arranged weekly. These two measures resulted in randomized above ground pairings between the two plant species, thus allowing us to exclude systematic effects of above ground interactions on root physiology and resistance.

### Analysis of root VOC profiles in the gap

To characterize the VOCs that accumulate in the gap between *T. officinale* and *C. stoebe*, we collected and analyzed VOCs using solid phase microextraction (SPME) and gas chromatography mass spectrometry (GC-MS). After seven weeks of transplantation, VOCs were collected from two randomly selected pots of each combination for one biological replicate (n = 4 per combination). An SPME fiber (coated with 100 μm polydimethylsiloxane; Supelco, Bellefonte, PA, USA) was inserted into the gap of a pot and exposed to VOCs for 60 min at room temperature and then transferred to another pot for 60 min for collection. Subsequently, the incubated fiber was immediately analyzed by GC-MS (Agilent 7820A GC interfaced with an Agilent 5977E MSD, Palo Alto, CA, USA) following previously established protocols (Huang et al., 2017). Briefly, the fiber was inserted into the injector port at 250°C and desorbed for 2 min. After insertion, the GC temperature program was 60 °C for 1 min, increased to 250 at 5°C min^−1^ and followed by 4 min at 250°C. The chromatograms were processed using default settings for spectral alignment and peak picking of PROGENESIS QI (Nonlinear Dynamics, Newcastle, UK). Features were assigned to individual compounds by retention time and peak shape matching and all VOCs were tentatively identified by the use of the NIST search 2.2 Mass Spectral Library (Gaithersburg, MD, USA) as well as retention time and spectral comparison with pure compounds as described (companion paper Gfeller et al., under review). During the experiment, the pots were watered every day and re-arranged every week.

### Contribution of (E)-β-caryophyllene to plant-plant interactions

(*E*)-β-caryophyllene is one of the major sesquiterpenes released by *C. stoebe* roots and is produced by the root-expressed terpene synthase CsTPS4 (companion paper Gfeller et al., under review). To test whether (*E*)-β-caryophyllene is sufficient to account for the increased growth of *M. melolontha* on *T. officinale* plants, we determined concentration of (*E*)-β-caryophyllene in the airgap between the rhizosphere of *C. stoebe* and *T. officinale* (see above) and then used corresponding synthetic doses to investigate its impact on the interaction between *T. officinale* and *M. melolontha*.

To check whether we can mimick the (*E*)-β-caryophyllene release of *C. stoebe* with a dispenser containing synthetic (*E*)-β-caryophyllene, we measured (*E*)-β-caryophyllene in the airgap of *T. officinale* plants growing with *C. stoebe* or *T. officinale* plants growing without *C. stoebe* but with an (*E*)-β-caryophyllene dispenser in the airgap (*n* = 16). Both plant species were seven-weeks old. Dispensers were constructed from 1.5 ml glass vials (VWR) that were pierced by a 1 ul micro-pipette (Drummond) and sealed with parafilm (Bemis). Dispensers were filled with with 100 ul (*E*)-β-caryophyllene (> 98.5%, GC, Sigma-Aldrich). This device allowed for constant release rates of (*E*)-β-caryophyllene. Two days after the dispensers were added, (*E*)-β-caryophyllene concentrations were determined by SPME-GC-MS as described above, resulting in eight biological replicates (two pooled setups per replicate).

To test the effect of (*E*)-β-caryophyllene on the interaction between *T. officinale* and *M. melolontha,* we conducted an experiment within which *T. officinale* plants were exposed to (1) control dispensers without neighboring plant, (2) (*E*)-β-caryophyllene dispensers without neighboring plant, and (3) control dispensers with *C. stoebe* as a neighboring plant (n = 12 per combination). The experimental setup was as described above. Seven weeks after the transplantation of *C. stoebe* and the addition of the dispensers, one pre-weighted and starved *M. melolontha* larva was added to the mesh cage in which the *T. officinale* plants were growing. After 18 days, all larvae were recovered from mesh cages and re-weighted. During the experiment, the dispensers were replaced every ten days and pots were re-arranged every week.

### Data analysis

All data analyses were performed with the statistical analysis software R 3.2.0 (R Foundation for Statistical Computing, Vienna, Austria) using ‘Car’, ‘Lme4’, ‘Lsmeans’, ‘Vegan’ And ‘Rvaidememoire’ packages (Bates, Mächler, Bolker, & Walker, 2015; Fox & Weisberg, 2011; Hervé, 2016; Lenth, 2016; Oksanen et al., 2016). Data was analyzed using One- or Two-way analyses of variance (ANOVAs). ANOVA assumptions were verified by inspecting residuals and variance. Multiple comparisons were carried out using least square mean post-hoc tests (LSM). *P*-values were corrected using the False Discovery Rate (FDR) method (Benjamini & Hochberg, 1995). Associations between variables were tested using Pearson’s Product-Moment correlations. To examine the overall differences in VOC profiles among different combinations, the relative abundance of the detected features was subjected to principal component analysis (PCA). Monte Carlo tests with 999 permutations were then used to test for significant differences between combinations.

## Results

### Neighbor identity determines VOC profiles in the rhizosphere

**Fig. 1.**
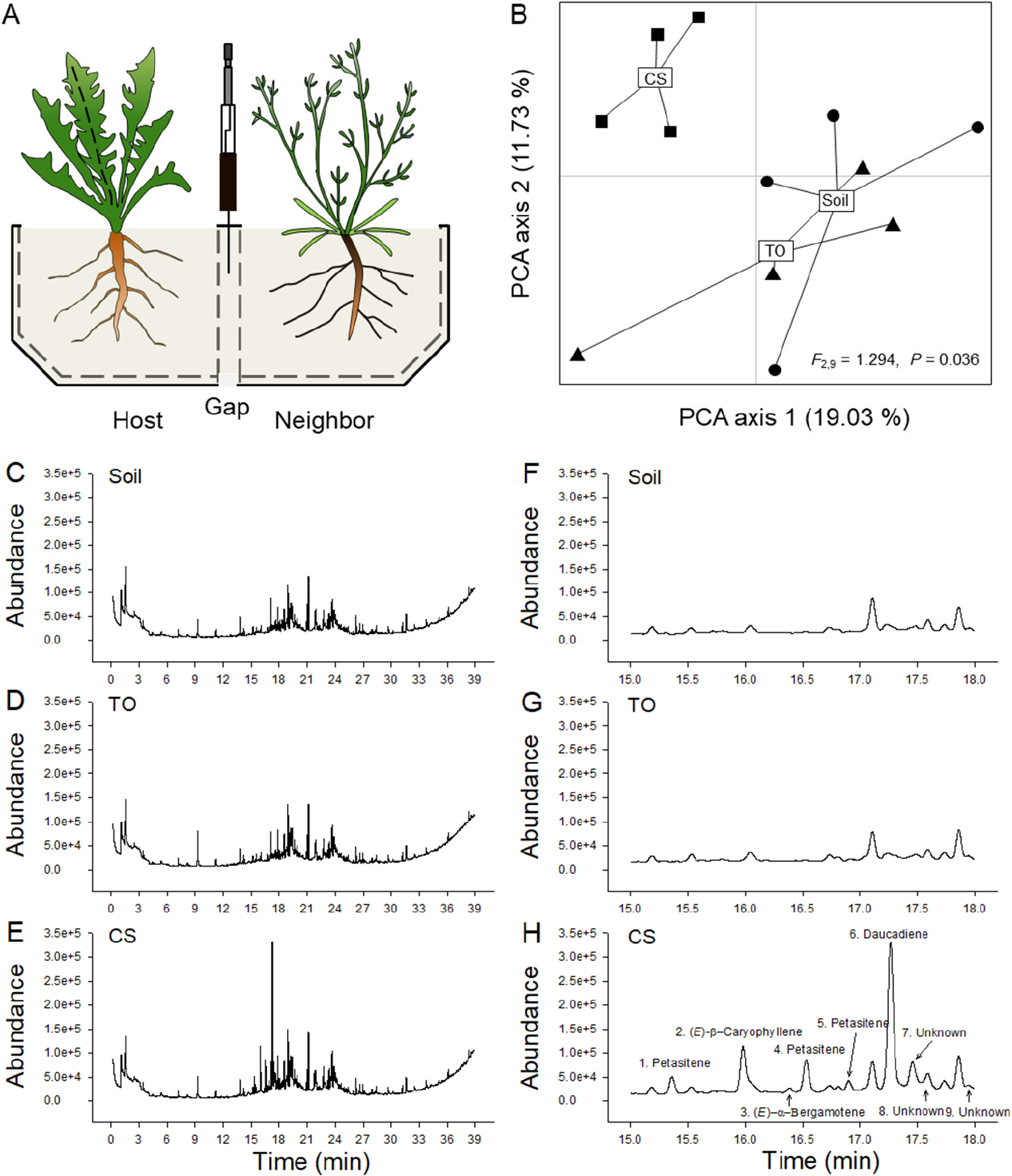
Sesquiterpene VOCs from *C. stoebe* diffuse through the rhizosphere. Experimental setup (A): *Taraxacum officinale* plants were grown in the vicinity of empty soil compartments (Soil), *T. officinale* plants (TO) or *Centaurea stoebe* (CS) plants, and volatiles were collected in the gap between the plants. The results of a principal component analysis of the VOC profiles in the gap are shown (B): The first two axes explained 19.03% and 11.73% of the total variation, respectively. Differences between treatments were determined by PCA. Data points represent biological replicates (n = 4). Circle, regular triangle and inverted triangle indicate neighbor identity, including Soil, TO and CS, respectively. Average abundance of GC-MS chromatograms of volatiles collected from gap between focal and neighboring plants from 0 to 39 min (C-E) and from 15 to 18 mins (F-H).

PCA analysis revealed that VOC profiles in the airgap between the *T. officinale* rhizosphere and the rhizospheres of the neighboring treatments differed significantly (*r*^*2*^ = 0.457, *P* = 0.009, Fig. 1). VOC profiles of *T. officinale* plants exposed to bare soil or *T. officinale* plants were indistinguishable (*P* = 0.516; Fig. 1B). By contrast, profiles were strongly altered by the presence of *C. stoebe* (*P* = 0.040, Fig. 1B). VOC profiles in the airgap between *T. officinale* and *C. stoebe* were dominated by sesquiterpenes that are released by *C. stoebe* roots (companion paper Gfeller et al., under review), including petasitenes, (*E*)-β-caryophyllene and daucadiene (peak area, *P* < 0.05, Fig. 1C-H).

### Root VOCs of C. stoebe increase M. melolontha growth on T. officinale

The growth of *M. melolontha* was similar on *T. officinale* plants that received below ground VOCs from bare soil or *T. officinale* neighbors (*P* = 0.791, Fig. 2B). By contrast, *M. melolontha* weight gain was significantly higher on *T. officinale* plants that were exposed to root VOCs of *C. stoebe* (*P* = 0.045, Fig. 2B). Thus, *C. stoebe* root VOCs increase *M. melolontha* growth on *T. officinale*.

**Fig. 2.**
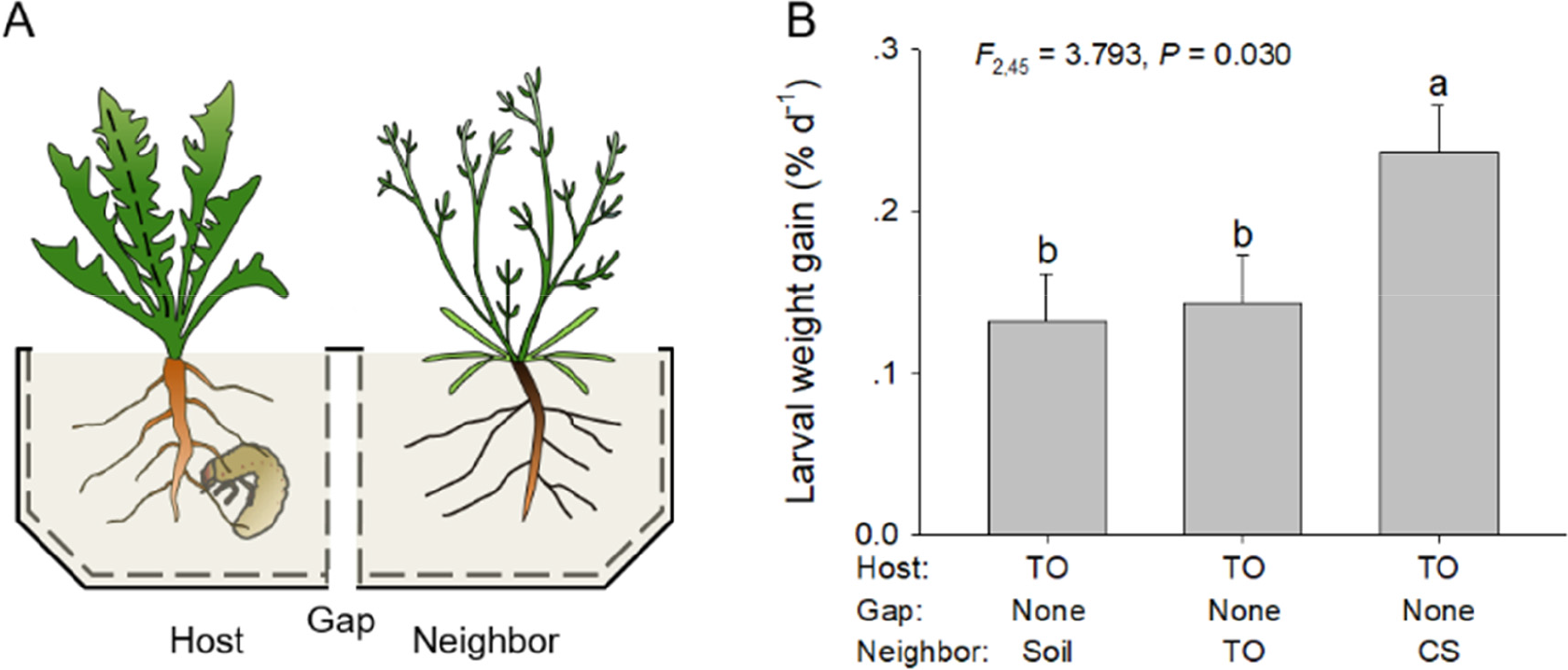
Root VOCs emitted by *C. stoebe* determine *Melolontha melolontha* performance. Experimental setup (A): Individual *Melolontha melolontha* larvae were allowed to feed on *Taraxacum officinale* plants growing in the vicinity of empty soil compartments (Soil), *T. officinale* (TO) or *Centaurea stoebe* (CS) for 18 days. Larval performance (B): Average larval weight gain was calculated as percentage increase in larval weight per day and is shown as mean ± 1 SE (n = 16). Differences between treatments were determined by One-way ANOVAs followed by post hoc multiple comparisons (different letters indicate *P* < 0.05, LSM).

### Root VOCs of C. stoebe change primary metabolites in T. offinicale roots

*Taraxacum officinale* root biomass was significantly affected by the different VOC exposure treatments (F_2,66_ = 5.571, *P* = 0.006), but not by *M. melolontha* attack (*F*_1,66_ = 1.761, *P* = 0.189) or the interaction (*F*_2,66_ = 0.253, *P* = 0.777). Root biomass was increased for plants exposed to *C. stoebe* root VOCs compared to plants that were exposed to *T. officinale* VOCs (Fig. 3A). Root VOC exposure also influenced the concentration of root primary and secondary metabolites (Fig. 3B-F). Total root protein concentrations were significantly affected by the VOC source (*F*_2,66_ = 50.383, *P* < 0.001), *M. melolontha* attack (*F*_1,66_ = 16.351, *P* < 0.001) and their interaction (*F*_2, 66_ = 12.506, *P* < 0.001). *Melolontha melolontha* attacked roots had higher protein levels upon exposure to *C. stoebe* (Fig. 3B). Root glucose levels were significantly affected by the VOC source (*F*_2,66_ = 6.841, *P* = 0.002), but not by *M. melolontha* attack (*F*_1,66_ = 0.023, *P* = 0.880) or their interaction (*F*_2,66_ = 3.118, *P* = 0.051). In the absence of *M. melolontha,* root glucose levels were higher in *C. stoebe* and *T. officinale* exposed plants compared to plants exposed to bare soil (Fig. 3C). Root fructose and sucrose were significantly affected by neighbor identify (fructose: *F*_2,66_ = 3.810, *P* = 0.027; sucrose: *F*_2,66_ = 4.595, *P* = 0.014) and *M. melolontha* attack (fructose: *F*_1,66_ = 10.346, *P* = 0.002; sucrose: *F*_1,66_ = 5.659, *P* = 0.020), but not by their interaction (fructose: *F*_2,66_ = 2.661, *P* = 0.077; sucrose: *F*_2,66_ = 0.699, *P* = 0.501) (Fig. 3D-E). Root sucrose levels were lower in *C. stoebe* exposed plants compared to *T. officinale* exposed plants in the absence of *M. melolontha* (Fig. 3E). The secondary metabolite TA-G was significantly decreased when *T. officinale* was attacked by *M. melolontha* larvae (*F*_1,66_ = 4.339, *P* = 0.041), but was not affected by the VOC source (*F*_2,66_ = 2.741, *P* = 0.072) or their interaction (*F*_2,66_ = 0.157, *P* = 0.855) (Fig. 3F). Thus, *T. officinale* plants respond to root VOCs from neighboring *C. stoebe* plants by increasing root growth and the abundance of primary metabolites. Root protein levels are also changed upon *C. stoebe* root VOC exposure, but this effect is only significant in combination with *M. melolontha* attack. Across treatments, *M. melolontha* larval weight gain was positively correlated with *T. officinale* root biomass (P = 0.007, *R*^2^ = 0.186, Fig. 3G) and soluble protein (*P* = 0.001, *R*^2^ = 0.272, Fig. 3H), but not significantly correlated with soluble sugars (Glucose, *P* = 0.255, *R*^2^ = 0.035; Fructose, *P* = 0.507, *R*^2^ = 0.010; Sucrose, *P* = 0.233, *R*^2^ = 0.053; Fig. 3I-K) or TA-G (*P* = 0.255, *R*^2^ = 0.034, Fig. 3L).

**Fig. 3.**
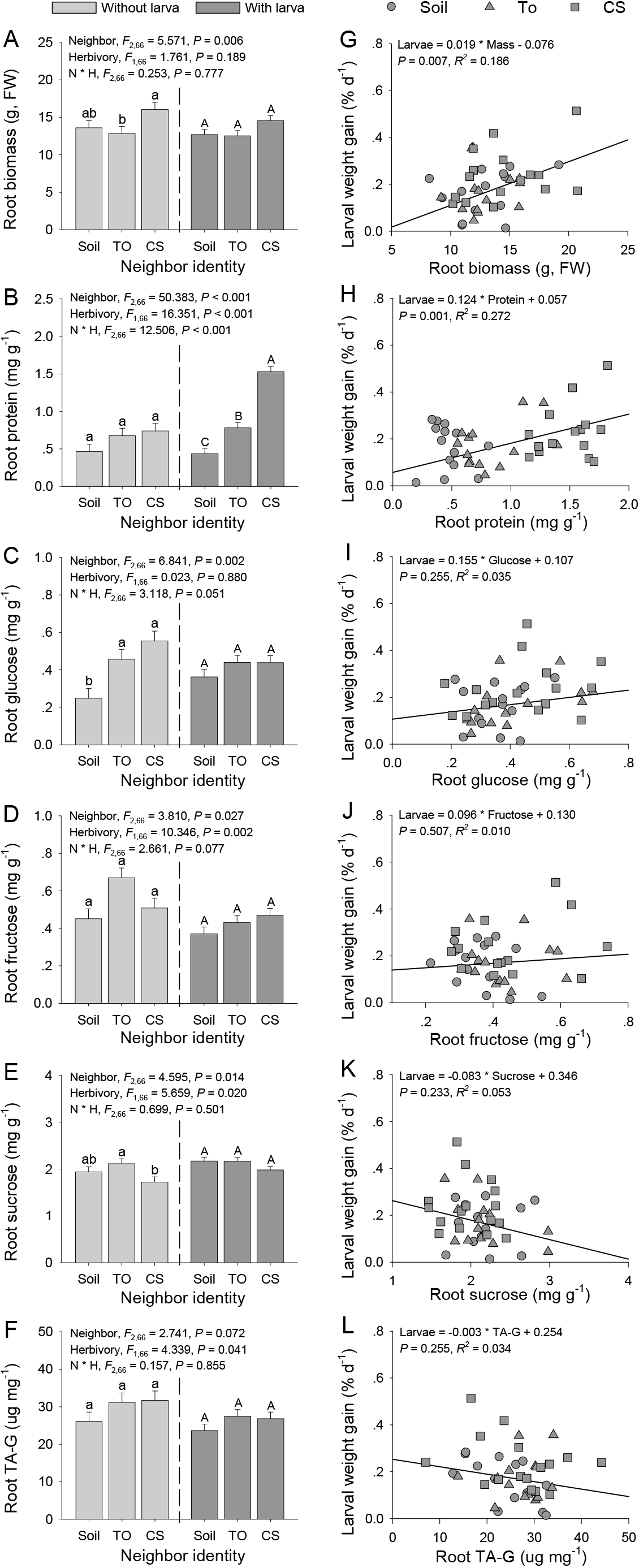
Root VOCs emitted by neighboring plant influence growth and chemistry of *Taraxacum officinale*. Root biomass (A), soluble protein (B), glucose (C), fructose (D), sucrose (E) and TA-G (F) of *T. officinale* growing in the vicinity of empty soil compartment (Soil), *T. officinale* (TO) or *Centaurea stoebe* (CS) are shown on the left. *T. officinale* plants were not attacked (light grey bars, n = 8) or attacked by *Melolontha melolontha* larvae (dark grey bars, n = 16). Values are means ± 1 SE. Differences between treatments were determined by Two-way ANOVAs followed by post hoc multiple comparisons (different letters in each herbivory group indicate *P* < 0.05, LSM). The relationships between larval weight gain and root biomass (G), soluble protein (H), glucose (I), fructose (J), sucrose (K) and TA-G (F) of *T. officinale* are shown on the right. Circle, regular triangle and inverted triangle indicate *T. officinale* growing in the vicinity of Soil, TO or CS, respectively. Regression equations, *P*-values and *R*^2^ values and are shown in the top of each figure.

### Synthetic (E)-β-caryophyllene partially mimics C. stoebe root VOC effects

The amount of (*E*)-β-caryophyllene that accumulated in the airgap supplied with a dispenser was similar to the emission of (*E*)-β-caryophyllene into the gap by *C. stoebe* (*t* = −0.302, *P* = 0.767, Fig. 4B). Similar to the previous experiment, the presence of *C. stoebe* increased *M. melolontha* weight gain compared to bare soil (Fig. 4C). *M. melolontha* growth in the presence of (*E*)-β-caryophyllene dispensers was intermediate and not statistically different from the control treatment or the *C. stoebe* treatment (Fig. 4C). Thus, (*E*)-β-caryophyllene partially mimics *C. stoebe* root VOC effects on *M. melolontha* growth on neighboring plants.

**Fig. 4.**
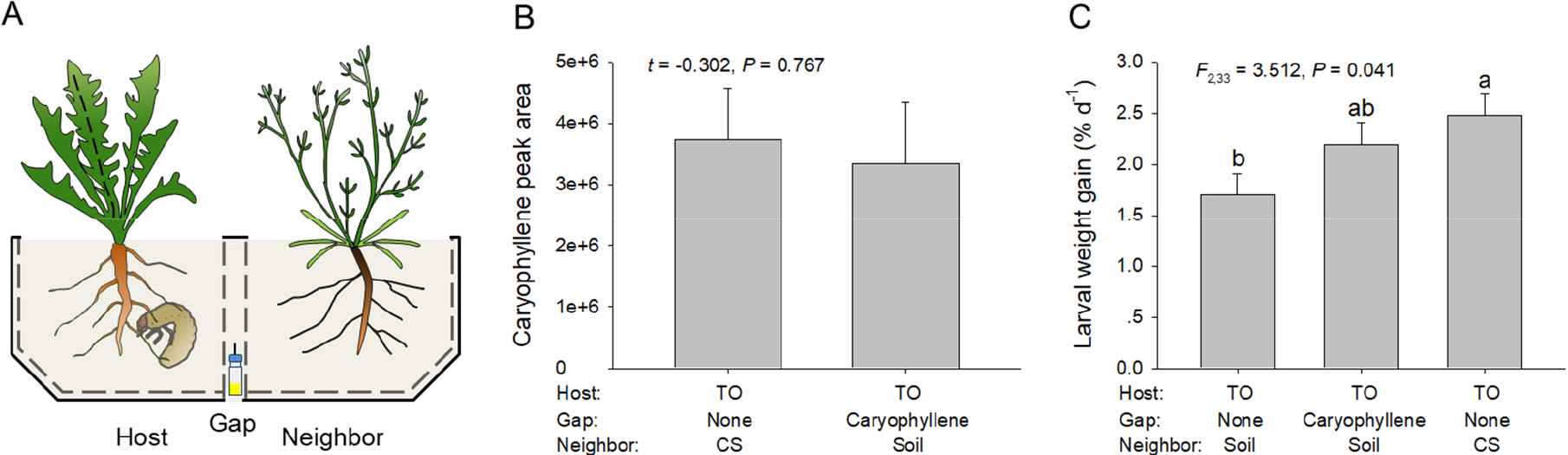
(*E*)-β-caryophyllene contributes to increased *Melolontha melolontha* growth on neighboring plants. Experimental setup (A): *Taraxacum officinale* plants were growing in the vicinity of empty soil compartment (Soil) or *Centaurea stoebe* (CS) and supplemented with or without synthetic (*E*)-β-caryophyllene in the gap. Physiological concentration of (*E*)-β-caryophyllene in gap (B): Control and (*E*)-β-caryophyllene dispensers were put in the gap for two days before measurements. Values were mean ± 1SE (n = 8). Differences between treatments were determined by independent sample *t*-tests. Impact of (*E*)-β-caryophyllene on *M. melolontha* larval growth (C): *Melolontha melolontha* larva was allowed to feed on *Taraxacum officinale* for 18 days. Values were mean ± 1SE (n = 12). Differences between treatments were determined by One-way ANOVA followed by post hoc multiple comparisons (different letters indicate *P* < 0.05, LSM).

## Discussion

Associational effects triggered by plant VOCs play important roles in determining plant-herbivore interactions in the field (Barbosa et al., 2009; Underwood, 2014). However, to date, most studies focused on above ground interactions through airborne signals, and most studies document that leaf VOCs trigger associational resistance in neighbors (Arimura et al., 2000; Engelberth et al., 2004; Frost et al., 2008; Erb et al., 2015; Pearse et al., 2013; Sugimoto et al., 2014). Our results show that root VOCs modulate plant-herbivore interactions and that VOCs may lead to associational susceptibility.

In an earlier study, we found that the presence of *C. stoebe* enhanced the performance of *M. melolontha* larvae feeding on *T. officinale* roots (Huang et al., 2018). In general, physical (e.g. light and contact), chemical (e.g. volatile and exudates) and biological (e.g. arbuscular mycorrhizal fungi) factors may trigger neighborhood effects and affect plant growth and defense (Babikova et al., 2013; Crepy & Casal, 2015; Erb et al., 2015; Hu et al., 2018b; Kong et al., 2018; Semchenko, Saar, & Lepik, 2014; Yang, Callaway, & Atwater, 2015). As *C. stoebe* constitutively releases large amounts of sesquiterpenes into the rhizosphere (companion paper Gfeller et al., under review), we hypothesized that root VOCs may be responsible for the plant-mediated changes in *M. melolontha* growth. Using an experimental setup that effectively randomizes above ground cues and eliminates root contact and the exchange of soluble exudates, we found that *C. stoebe* root volatiles diffuse through the rhizosphere and are sufficient to increase the growth of *M. melolontha* on neighboring *T. officinale*. Thus, this study provides experimental evidence that root VOCs play an important role in below ground associational effects impacting plant-herbivore interactions.

Associational effects elicited by plant VOCs can be the result of chemical changes of receiver plants (Engelberth et al., 2004; Erb et al., 2015; Huang et al., 2018; Sugimoto et al., 2014). In our earlier work, we excluded the possibility that *M. melolontha* is directly affected by *C. stoebe* root VOCs or exudates, suggesting that *C. stoebe* increases *M. melolontha* growth through plant-mediated effects. In line with this hypothesis, we demonstrate here that growth and primary metabolism of *T. officinale* roots changes upon exposure to root VOCs of *C. stoebe*. Some of these effects are even stronger when the plants are attacked by *M. melolontha*, suggesting an interaction between root VOC exposure and herbivory. For instance, exposure to *C. stoebe* root VOCs increases root protein content and root growth of *T. officinale* plants. Both parameters are positively correlated with larval performance, indicating that *M. melolontha* growth may be stimulated by enhanced root growth and nutrient levels. Previous studies demonstrated that secondary metabolites such as TA-G protect *T. officinale* against *M. melolontha* (Bont et al., 2017; Huber et al., 2016a; Huber et al., 2016b). We found on clear effects of *C. stoebe* VOCs on root TA-G concentrations, implying that *C. stoebe* VOCs do not act by suppressing this plant defense.

The identification of bioactive VOCs from plant-derived blends remains an important bottleneck in chemical ecology. We show that *C. stoebe* releases a complex blend of sesquiterpenes as well as other minor unidentified VOCs from its roots (companion paper Gfeller et al., under review), all of which may be associated with the observed effects on *M. melolontha* growth. Here, we tested whether (*E*)-β-caryophyllene, one of the major sesquiterpenes emitted by *C. stoebe*, is sufficient to increase the growth of *M. melolontha* on *T. officinale* in comparison with the full VOC blend of *C. stoebe.* (*E*)-β-caryophyllene is a widespread sesquiterpene in nature that can influence the physiology and behavior of fungi, nematodes and insects (Fantaye, Köpke, Gershenzon, & Degenhardt, 2015; Rasmann et al., 2005; Robert et al., 2013) and may act as an antioxidant in plants (Palmer-Young, Veit, Gershenzon, & Schuman, 2015). We demonstrate that (*E*)-β-caryophyllene exposure leads to *M. melolontha* growth that is intermediate between non-exposed and *C. stoebe* exposed *T. officinale* plants, suggesting that it can partially account for the VOC effects of *C. stoebe*. We propose that other sesquiterpenes emitted by *C. stoebe* such as daucadiene and petasitene, may also contribute to enhanced *M. melolonta* growth. More work is needed to test this hypothesis. The identification of TPSs that are likely responsible for sesquiterpene production in *C. stoebe* (companion paper Gfeller et al., under review) represents a first step towards the manipulation and functional assessment of *C. stoebe* root VOCs *in vivo*.

VOCs of neighboring plants are well known to increase defenses and resistance of neighboring plants (Arimura et al., 2000; Erb et al., 2015; Sugimoto et al., 2014), and only few documented examples exist where VOCs decrease the resistance of neighboring plants (Li & Blande, 2015, Erb, 2018). From the perspective of the sender, inducing susceptibility to herbivores in neighboring plants may be an advantage, as it may reduce their competitiveness. VOC-induced susceptibility may thus be a form of plant offense. However, several caveats need to be considered. First, many herbivores are mobile, and increasing herbivore growth on neighboring plants may lead to accelerated migration to the sender plant. Second, herbivore growth, as measured here, is not synonymous with plant damage and may be the result of an increase in performance of the receiver plant, in which case their competitiveness would not be reduced, and the benefit for the emitter would be less evident (Erb, 2018a; Veyrat, Robert, Turlings, & Erb, 2016). Third, the benefits of inducing susceptibility in neighboring plants may be offset in the absence of herbivores. Indeed, we show that *C. stoebe* VOCs can increase germination and growth of heterospecific neighboring plants in the absence of herbivores (companion paper Gfeller et al., under review). Therefore, more research is needed to understand the evolutionary and ecological context of the present findings.

In conclusion, the present study shows that root VOCs can influence plant-herbivore interactions on neighboring plants through plant-mediated effects. Thus, associational effects mediated by below ground VOCs need to be included into models on plant community ecology.

## Acknowledgements

We are grateful to Noelle Schenk for insect rearing and the gardeners of the IPS for plant cultivation. This study was supported by the Swiss National Science Foundation (grants nos’. 153517 and 157884 to M.E.), the European Commission (MC-CIG no. 629134 to M.E., MC-IEF no. 704334 to W.H.), and the National Natural Science Foundation of China (31470447 and 31822007to W.H.). The authors declare that they have no conflict of interest.

## Author contributions

W.H. and M.E. designed the experiments. W.H. carried out greenhouse research. W.H., V.G. and M.E. performed chemical analyses, analyzed data and wrote the manuscript.

## Data accessibility

Raw data associated with this study can be downloaded from Dryad [to be inserted at a later stage]

